# Identification of reduced host transcriptomic signatures for tuberculosis and digital PCR-based validation and quantification

**DOI:** 10.1101/583674

**Authors:** Harriet D. Gliddon, Myrsini Kaforou, Mary Alikian, Dominic Habgood-Coote, Chenxi Zhou, Tolu Oni, Suzanne T. Anderson, Andrew J. Brent, Amelia C. Crampin, Brian Eley, Florian Kern, Paul R. Langford, Tom H. M. Ottenhoff, Martin L. Hibberd, Neil French, Victoria J. Wright, Hazel M. Dockrell, Lachlan J. Coin, Robert J. Wilkinson, Michael Levin, on behalf of the ILULU Consortium

## Abstract

Recently, host whole blood gene expression signatures have been identified for diagnosis of tuberculosis (TB). Absolute quantification of the concentrations of signature transcripts in blood have not been reported, but would facilitate the development of diagnostic tests.

To identify minimal transcript signatures, we applied a novel transcript selection procedure to microarray data from African adults comprising 536 patients with TB, other diseases (OD) and latent TB (LTBI), divided into training and test sets. Signatures were validated using reverse transcriptase (RT) - digital PCR (dPCR).

A four-transcript signature (*GBP6*, *TMCC1*, *PRDM1*, *ARG1*) measured using RT-dPCR distinguished TB patients from those with OD (area under the curve (AUC) 93.8% (CI_95%_ 82.2 – 100%). A three-transcript signature (*FCGR1A, ZNF296, C1QB*) differentiated TB from LTBI (AUC 97.3%, CI_95%_: 93.3 – 100%), regardless of HIV.

These signatures have been validated across platforms and across samples offering strong, quantitative support for their use as diagnostic biomarkers for TB.

## Introduction

Despite over a century’s research effort to identify new diagnostic tools we still lack diagnostic tests for tuberculosis (TB) that are sensitive, affordable and robust. The majority of TB diagnostics are based on identifying the pathogen in sputum, by microscopy, culture or PCR. However, current methods fail to identify the pathogen in a significant proportion of cases, either due to inadequacies in sputum collection, paucibacillary disease, HIV infection or in patients with extrapulmonary forms [1]. As a result the World Health Organization (WHO) estimates that 37% of TB cases go unreported or undiagnosed [2]. Given the problems associated with using sputum as a clinical sample, the WHO and the Foundation for Innovative New Diagnostics published a target product profile (TPP) for a non-sputum biomarker test in 2014 [3]. This specified the seven proposed key characteristics of a rapid biomarker-based non-sputum-based test for detecting TB including minimal and optimal sensitivity and specificity of such a test and also discussed sample accessibility, time to result, maintenance and cost.

Recent years have seen a revival of host-response based diagnostics in the focus of diagnostic tests for infectious diseases that detect evidence of a host immune response to an infection, which is advantageous when there are very low numbers of the pathogen in the body or when pathogens colonise inaccessible sites. A number of disease specific ‘omic’ signatures have been identified, facilitated by advances in technology to analyse the genome, transcriptome, epigenome, lipidome, metabolome and proteome in a high-throughput and quantitative manner [4]. As well as improving our understanding of the pathogenesis of a range of infectious diseases, these signatures have the potential to be used as diagnostic biomarkers.

Gene expression studies have significantly enhanced our knowledge of the roles of various components of the immune system in TB disease [5] [6] [7]. A number of gene expression signatures have been published that can distinguish TB from healthy controls (HCs) [5] [8] [9] and correlate with disease progression [8] [10] [11]. These could serve as important indicators of disease progression from latent TB infection (LTBI) to TB, and therefore aid the targeting of appropriate antibiotic treatment [12].

The most clinically important need is for biomarkers to distinguish TB from the range of other conditions with similar clinical presentation. TB shares symptoms, clinical signs and laboratory findings with many other diseases (OD), including a wide range of infectious, inflammatory and malignant conditions, such as pneumonia or other HIV-associated opportunistic infections. Distinguishing between TB and OD is particularly important in patients living with HIV, because extrapulmonary TB is more common in these patients [13] [14] such that most sputum-based tests are poorly sensitive, and HIV-associated malignancies or opportunistic infections can have similar clinical presentations. However, the majority of TB gene expression studies published to date have compared TB cohorts to HCs, LTBI or patients with OD, mostly in the absence of HIV infection.

A previous study (Kaforou et al., [9]) addressed these issues by studying patients with symptoms suggestive of TB in Malawi and South Africa (including both HIV-infected and uninfected persons) and classifying them as TB, LTBI or OD. Blood gene expression signatures were identified using genome-wide microarrays that distinguished TB from OD and LTBI [9]. A 44-transcript signature was found to distinguish TB from OD with sensitivity of 93% (CI_95%_ 83-100) and specificity of 88% (CI_95%_ 74-97). A 27-transcript signature distinguished TB from LTBI with sensitivity of 95% (CI_95%_ 87-100) and specificity of 90% (CI_95%_ 80-97). These signatures showed only slightly reduced accuracy in HIV-coinfected individuals [9].

Further reduction in the number of transcripts comprising these gene expression signatures makes their use as diagnostic markers more feasible for clinical translation, particularly at the point-of-care and in resource-limited settings [15]. This has been the subject of significant research effort and a number of bioinformatics approaches have been employed. Sweeney et al. identified a three-gene signature for TB, comprised of *GBP5*, *DUSP3*, and *KLF2* in a meta-analysis of publicly available gene expression microarray data [16]. Maertzdorf et al. used random forest models and confidence interval decision trees to identify a four-transcript signature comprising *GBP1*, *IFITM3*, *P2RY14* and *ID3*, that distinguished between TB and HC, regardless of HIV infection status [17]. Other recent studies identified minimal gene expression signatures in populations from high-endemic countries that predict progression from latent infection to active TB disease with accuracy, excluding cases with HIV co-infection [18] [19].

Quantification of individual TB gene expression signature transcripts would be useful to determine the limits of detection required for diagnostic tests based on these signatures. The established method of choice for performing absolute quantification of nucleic acids is quantitative PCR (qPCR), where amplicon generation is measured in real time and related back to the starting concentration of template. While RNA-seq has emerged as a powerful technique for investigating RNA species within a given sample, it can only provide *relative* quantification of RNA species [20]. In recent years, digital PCR (dPCR) has emerged as a promising alternative to qPCR. dPCR is a useful method for quickly and efficiently providing absolute quantification of individual mRNA species and has been shown to be more reproducible and less prone to inhibition than qPCR [21] [22]. The high precision offered by dPCR makes it ideally suited to the detection of rare point mutations and the accurate detection of low microbial loads, among other applications [23] [24] [25].

We hypothesised that we could further reduce the number of transcripts comprising the previously reported signatures distinguishing TB from OD and LTBI (Kaforou et al. [9]) using novel feature selection algorithms applied to microarray data, and that reverse transcription-dPCR (RT-dPCR) could be used to quantify the concentrations of individual gene transcripts in purified RNA from whole blood. We postulated that this cross-sample, cross-platform, cross-population study will aid the advance of the TB transcriptomics field towards developing and establishing the use of host transcriptomics for TB diagnosis.

## Materials and methods

### Ethics statement

The study was approved by the Human Research Ethics Committee of the University of Cape Town, South Africa (HREC012/2007), the National Health Sciences Research Committee, Malawi (NHSRC/447), and the Ethics Committee of the London School of Hygiene and Tropical Medicine (5212). Written information was provided by trained local health workers in local languages and all patients provided written consent.

### Analysis of gene expression microarray data and identification of reduced signatures using the Forward Selection-Partial Least Squares (FS-PLS) method

The patient cohorts recruited in South Africa and Malawi for the original prospective cohort microarray study were fully described previously, including the diagnostic procedures and patient assignment as TB, OD or LTBI [8]. In addition, the whole-blood genome-wide expression measured in this cohort was reported [8], and made publicly available at NCBI’s Gene Expression Omnibus, accessible through GEO Series accession number GSE37250. The microarray data was pre-processed as described in [8]. Data from the processed and normalised expression set were split randomly into training and test set (80-20 split). FS-PLS [26] [27] was employed in order to generate smaller gene expression signatures. FS-PLS is an iterative forward selection algorithm which at each step selects the most strongly associated variable after projecting the data matrix into a space orthogonal to all the variables previously selected. It combines the dimensionality reduction strength of PLS and the model simplicity and interpretability of FS regression. The classificatory performance of the signatures was evaluated in the test set using the disease risk score method (DRS), as in [9]. The derived signatures were further validated in two publicly available gene expression studies [8] [5] (Supplementary Methods). The FS-PLS code is available for download and use [27].

### Power calculations for RT-dPCR study size

For the retrospective RT-dPCR study, as the discrimination using the DRS had a binary outcome and followed a binomial distribution, in order to achieve a statistic significance level of 0.05, and assuming the dPCR sensitivity to be at least 75% for patient classification, we used 40 samples for each comparison (TB vs OD and TB vs LTBI) to assess the performance of each signature, with equal numbers of samples for each group (n_TB_=20, n_OD_=20, n_LTBI_=20) (S1, S2 Table). Samples were chosen at random from a microarray test patient cohort for TB vs OD, stratified for HIV status and country of origin, which had not been used to derive the signature. An additional 10 LTBI HIV-infected and 10 LTBI HIV-uninfected samples from the test microarray cohort were analysed.

### Patient characteristics for RNA samples used in the RT-dPCR

Patient recruitment was conducted in two highly contrasting study sites in Cape Town, South Africa and Karonga District, Northern Malawi. Patients were classified as having active TB disease only upon culture confirmation. Patients were deemed to have OD if they presented with symptoms that might suggest the possibility of TB disease, but for whom an alternative diagnosis was found and TB treatment was not administered. These patients were followed up 26 weeks post diagnosis to confirm they remained TB-free. Healthy LTBI controls were classified according to the results of interferon-gamma release assay (IGRA) and tuberculin skin test (TST) investigations [9].

### RNA purification from whole blood and storage

2.5 mL whole blood was collected at the time of recruitment (before or within 24 hours of commencing TB treatment in suspected patients) in PAXgene blood RNA tubes (PreAnalytiX), frozen within three hours of collection, and later extracted using PAXgene blood RNA kits (PreAnalytiX). RNA was shipped frozen and stored at −80 °C.

### Assessment of RNA purity and integrity

Before proceeding with reverse transcription, the RNA quality of the samples was assessed using an Agilent 2100 Bioanalyzer (Agilent Technologies, Santa Clara, CA, USA).

### Reverse Transcription of purified RNA from whole blood

RNA concentration was measured using a NanoDrop 2000c (Thermo Scientific) and 500 ng was used for the reverse transcription reaction in a total volume of 10 µL nuclease-free H_2_O. RT was performed in one batch using the High-Capacity cDNA RT Kit (Applied Biosystems) according to the manufacturer’s instructions. The cycle was 25 °C for 10 minutes, 37 °C for 120 minutes, 85 °C for 5 minutes, followed by a hold at 4 °C. cDNA samples were stored at −20 °C for fewer than six months before use.

### dPCR using the QuantStudio^TM^ platform

Up to 5 μL of RT product was added to 7.5 μL QuantStudio 3D Digital PCR Master Mix (Thermo Fisher Scientific), 0.75 μL of TaqMan Assay (20X) (Thermo Fisher Scientific) (see S3 Table) and the volume made up to 15 μL using nuclease-free H_2_O. At least one no template control was used for each TaqMan assay on each PCR run. The reaction mix was applied to each QuantStudio 3D Digital PCR 20K Chip (Applied Biosystems) according to the manufacturer’s instructions. The dPCR was run on a GeneAmp PCR System 9700 (Applied Biosystems) with a cycle of 10 minutes at 96 °C, followed by 39 cycles of 60 °C for 60 seconds and 98 °C for 30 seconds, followed by two minutes at 60 °C before holding at 10 °C. Chips were read, and absolute quantification (copies per µL) determined using the QuantStudio 3D Digital PCR Instrument (Thermo Fisher Scientific).

### Data analysis RT-dPCR

Data was exported and analysed using QuantStudio 3D AnalysisSuite Cloud Software Version 3.0.3 (Thermo Fisher Scientific). The quantification algorithm selected was Poisson. The software assesses whether the data on a chip is reliable based upon loading, signal, and noise characteristics and displays quality indicators for each chip. Any chip that gave a precision value of >10% was deemed to have failed and was repeated. Similarly, if the negative and positive wells did not separate into distinct populations, the sample and probe combination was repeated. This failure to separate into two populations could be caused by the chips leaking, evaporation or a loading issue of the sample onto the chip. This methodology is further explained in the supporting information (S1 Fig) and all dilutions, FAM call thresholds and lambda values are given in S4 Table, in accordance with the MIQE guidelines [21]. The output given by the QuantStudio software is in copies/µL. This value was then corrected according to the dilution of cDNA used for the dPCR in order to determine the absolute concentration of a given transcript in purified RNA samples (S3 Fig). RT-dPCR derived copies per µL values are reported. The DRS method was used to classify patients on the basis of log_2_ (copies per μL).

### Statistical Analysis

The datasets were analysed in ‘R’ Language and Environment for Statistical Computing version 3.4.1 [28] [29]. In order to evaluate the performance of the DRS as a binary classifier, the area under the curve (AUC) for a receiver operating characteristic (ROC) curve was calculated, as well as the sensitivity and specificity using pROC [28]. The calculation of the confidence intervals (CI) for the AUC was based on the DeLong method [30], an asymptotically exact method to evaluate the uncertainty of an AUC, except for the one case that AUC=100%, where we used a smoothed ROC followed by DeLong for the calculation of the lower 95% bound. For each data set we report the point estimate for sensitivity as the closest value >90% (as specified in the WHO TPP) and the corresponding specificity.

## Results

### Discovery and Validation of small signatures from microarray data using FS-PLS and DRS

In order to derive reduced gene expression signatures with diagnostic potential, the variable selection method, FS-PLS, was applied to the previously published microarray data (80% training set) (n=293 for TB vs OD, n=285 for TB vs LTBI including HIV co-infected cases), tested in the test set (n=76 TB vs OD and TB vs LTBI including HIV co-infected cases), and further validated in two other publicly available studies (Fig 1) [5] [8]. Using the FSPLS method we identified a signature comprising four transcripts for TB/OD in the training set and a signature comprising three transcripts for TB/LTBI; the signatures are detailed in Tables 1a and 1b, respectively. The TB/OD FS-PLS signature using the DRS had an AUC of 93.9% CI_95%_ (88.4 – 99.4%) in the 20% test set, which had not been used for discovery (Fig 2), sensitivity of 90.5 CI_95%_ (77.4 - 97.3) and specificity of 82.4% CI_95%_ (65.5 – 93.2), with confidence intervals overlapping with the previously identified the 44-transcript elastic net signature for TB/OD [9]. The TB/LTBI FS-PLS signature using the DRS had an AUC of 95.4% (CI_95%_ 91.2-99.6%) in the 20% test set, which had not been used for discovery, sensitivity of 91.9 CI_95%_ (78.1 – 98.3) and specificity of 84.6% CI_95%_ (69.5 - 94.1), with confidence intervals overlapping with the previously reported 27-transcript elastic net signature [9] (Fig 2 and S5 Table).

**Table 1.**
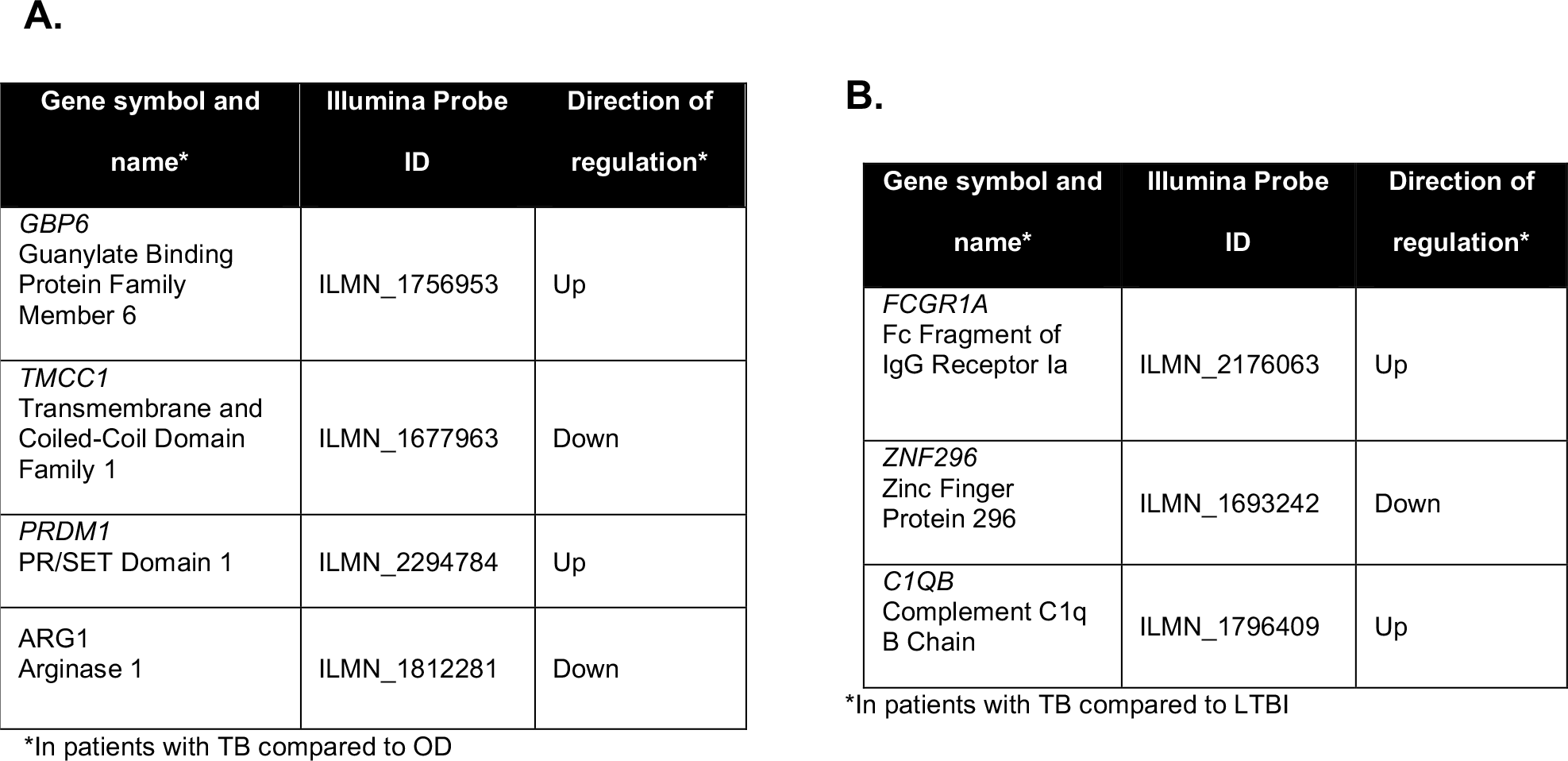
FS-PLS signatures for (a) TB/LTBI and (b) TB/OD. They were taken forward for further characterisation and validation using dPCR, and subsequently development as diagnostic biomarkers for TB. *According to HGNC.

**Fig 1.**
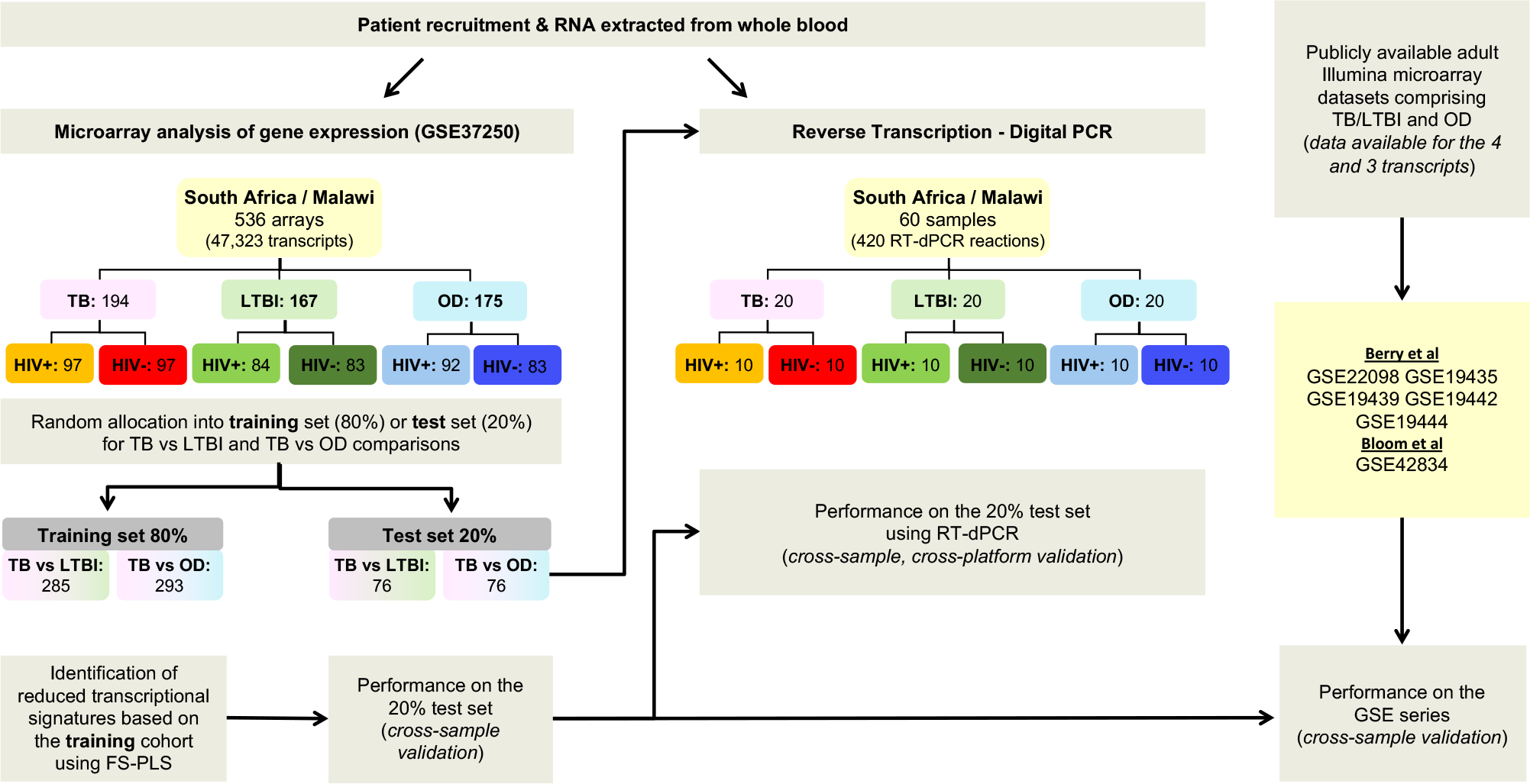
Workflow. Identification of small signatures for TB/LTBI and TB/OD from microarray data using (FS-PLS), followed by classification performance in a separate test set and finally, validation using RT-dPCR.

**Fig 2.**
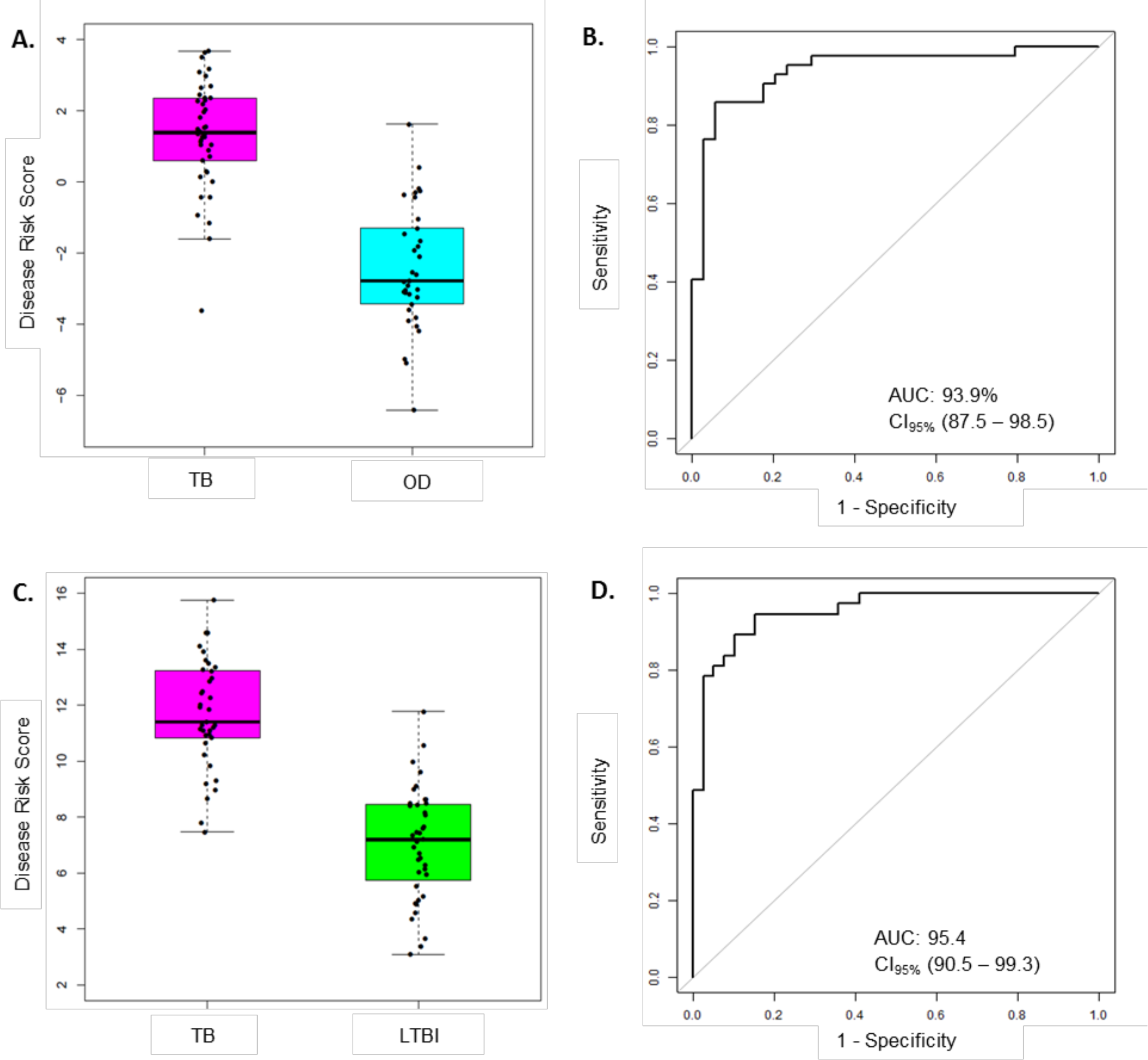
Classification performance of the FS-PLS-derived four-transcript signatures for TB/OD and three-transcript for TB/LTBI using microarray gene expression data (only test dataset shown). (A) Box plots of DRS and (B) ROC curve based on the TB/OD FS-PLS signature applied to the combined HIV +/− TB and OD SA/Malawi cohorts. (C) Box plots of DRS and (D) ROC curve based on the TB/LTBI FS-PLS signature applied to the combined HIV +/− TB and LTBI SA/Malawi cohorts.

### Validation of the FS-PLS TB/OD and TB/LTBI signatures in external datasets

In order to further validate the performance of the DRS based on the TB/OD four transcript and TB/LTBI three transcript signature, we employed the whole blood expression datasets of Berry et al. [5] and Bloom et al. [8] (GEO: GSE19491, GSE42834) as validation cohorts. The cohorts comprised HIV-uninfected individuals; TB, LTBI, and OD including pneumonia, lung cancer, Still’s disease, adult and pediatric Systemic Lupus Erythematosus (ASLE, PSLE), *Staphylococcus*, and *Streptococcus*. (Table 2). In comparison to the Maertzdorf et al. [17], our four-transcript signature had higher point estimates for the TB vs OD Berry et al. [5] datasets. In the Maertzdorf et al., an AUC of 71% (63– 71%) is reported for the Maertzdorf four-transcript signature when applied to the our TB/OD HIV-infected and HIV-uninfected microarray dataset [9],

**Table 2.**
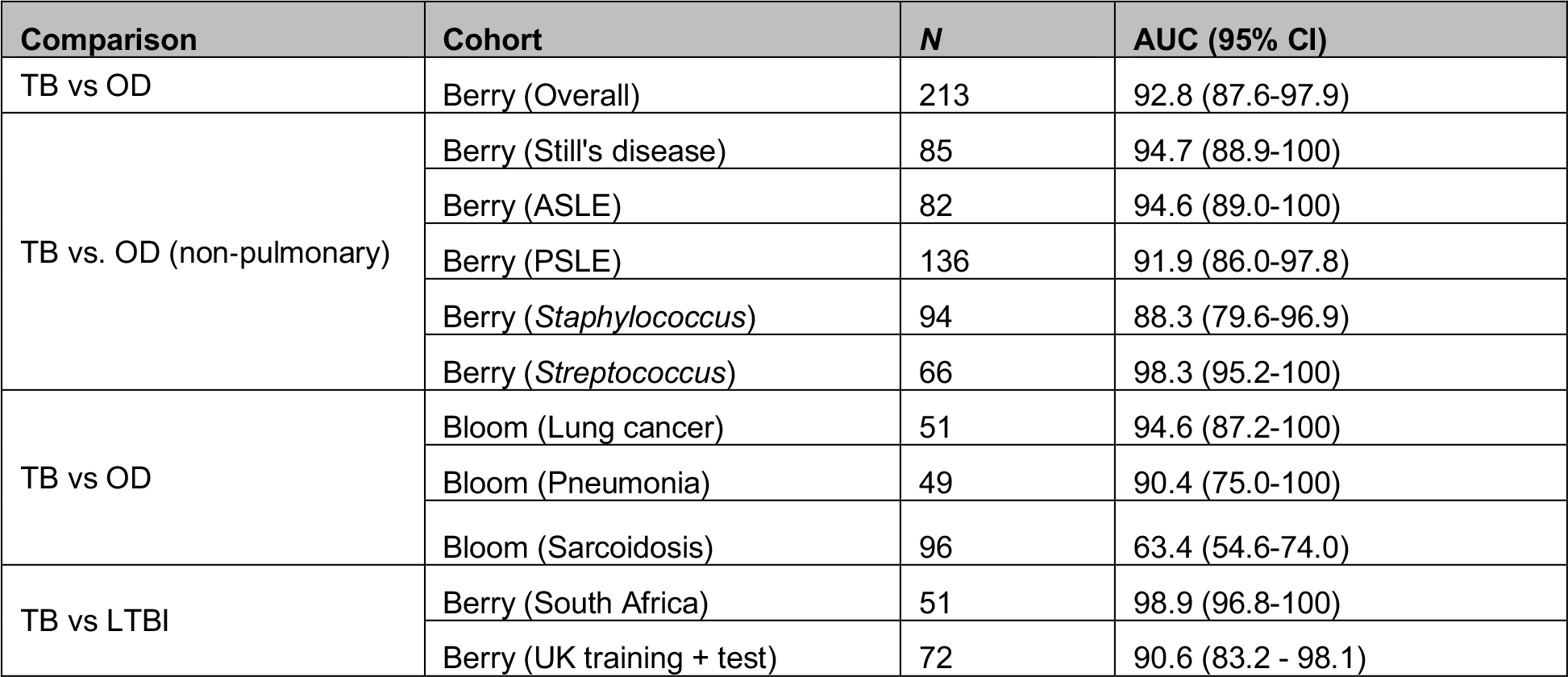
Performance of FS-PLS signatures in classifying TB and OD, or TB and LTBI, in other publicly available microarray datasets.

### Clinical characteristics of cohorts used in RT-dPCR analysis

The clinical characteristics of each disease cohort used for RT-dPCR analysis with the TB/LTBI signature genes and the TB/OD signature genes are shown in Table 3a and 3b, respectively. The mean age, body mass index (BMI) and TST induration are shown. The clinical diagnoses of the OD cohort are listed in S2 Table. The range of diagnoses among this cohort is representative of the variety of conditions that have similar clinical presentations to TB.

**Table 3.** Clinical characteristics of patients used for the RT-dPCR analysis of the (A) TB/OD signature and (B) the TB/LTBI signature.

| Group | TB HIV+ |  | TB HIV- |  | OD HIV+ |  | OD HIV- |  |
| --- | --- | --- | --- | --- | --- | --- | --- | --- |
| Location | SA | MLW | SA | MLW | SA | MLW | SA | MLW |
| Number | 6 | 4 | 5 | 5 | 6 | 4 | 6 | 4 |
| Age (years) mean (range) | 27.9<br>(26.6-33.9) | 35.4<br>(29.2-48.0) | 29.7<br>(19.4-60.7) | 23.5<br>(18.1-48.9) | 33.8<br>(24.9-36.5) | 41.3<br>(33.1-53.1) | 48.8<br>(19.1-68.1) | 34.1<br>(21.7-53.9) |
| Sex (male, %) | 50 | 50 | 60 | 40 | 33.3 | 25 | 50 | 0 |
| BMI (kg/m<sup>2</sup>) mean (range) | 20.5<br>(16.8-22.9) | 18.0<br>(16.9-18.8) | 19.3<br>(15.3-24.2) | 18.6<br>(16.5-20.2) | 26.7<br>(21.3-39.3) | 21.7<sup>a</sup><br>(20.3-23.2) | 21.6<sup>b</sup><br>(10.0-33.0) | 19.0<br>(17.2-20.8) |
| CD4 count (mm<sup>3</sup>) mean (range) | 266.8<br>(34.0-646.0) | 155.9<br>(29.2-337.1) | NA | NA | 336.2<br>(148-838) | 125.7<br>(11.3-204.8) | NA | NA |
| Anti-retroviral therapy | 0<br>(0%) | 0<br>(0%) | NA | NA | 3<br>(50%) | 3<br>(75%) | NA | NA |

<sup>a</sup> One missing value; <sup>b</sup> One missing value; NA Not applicable

| Group | TB HIV+ |  | TB HIV- |  | LTBI HIV+ |  | LTBI HIV- |  |
| --- | --- | --- | --- | --- | --- | --- | --- | --- |
| Location | SA | MLW | SA | MLW | SA | MLW | SA | MLW |
| Number | 6 | 4 | 5 | 5 | 5 | 5 | 5 | 5 |
| Age (years) mean (range) | 30.8<br>(26.3-38.8) | 34.8<br>(29.2-43.7) | 36.7<br>(29.7-60.7) | 48.0<br>(29.0-59.8) | 30.9<br>(22.5-38.2) | 42.9<br>(28.2-53.2) | 21.7<br>(18.9-23.9) | 42.1<br>(28.2-59.2) |
| Sex (male, %) | 50 | 50 | 60 | 40 | 0 | 80 | 20 | 20 |
| BMI (kg/m<sup>2</sup>) mean (range) | 21.9<br>(16.8-28.7) | 18.9<br>(16.9-22.4) | 21.0<br>(15.3-29.4) | 18.7<br>(14.3-21.4) | 26.8<br>(24.1-31.1) | 20.1<br>(16.5-25.7) | 23.5<br>(20.7-29.4) | 20.2<br>(17.7-23.4) |
| CD4 count (mm<sup>3</sup>) mean (range) | 215.3<br>(37.0-345.0) | 239.9<br>(29.2-337.1) | NA | NA | 630.0<sup>a</sup><br>(454.0-948.0) | 378.0<br>(227.0-545.5) | NA | NA |
| Anti-retroviral therapy | 0<br>(0%) | 0<br>(0%) | NA | NA | 0<br>(0%) | 0<br>(0%) | NA | NA |
| TST induration (mm) mean (range) | ND | ND | ND | ND | 24 (10-50) | 24.4 (22-28) | 17 (10-20) | 16.2 (11-21) |

<sup>a</sup> One missing value; ND Not done; NA Not applicable

### Absolute quantification of genes comprising the four-transcript FS-PLS signature for TB/OD (cross-platform, cross-sample validation)

Fig 3A shows the concentration (in copies per μL) of each of the transcripts comprising the FS-PLS signature for TB/OD in purified RNA from whole blood. *GBP6* transcript levels are higher in TB patients, compared to those with OD. The opposite case is observed for the *ARG1* transcript, which is more abundant in patients with OD compared to TB. For *TMCC1* and *PRDM1*, there is more overlap between concentration values of TB and OD patients. All four of these genes were identified in the 44 gene expression signature for TB/OD [9]. The original concentration (in copies per μL) for the samples stratified by HIV status is shown in S2 Fig.

**Fig 3.**
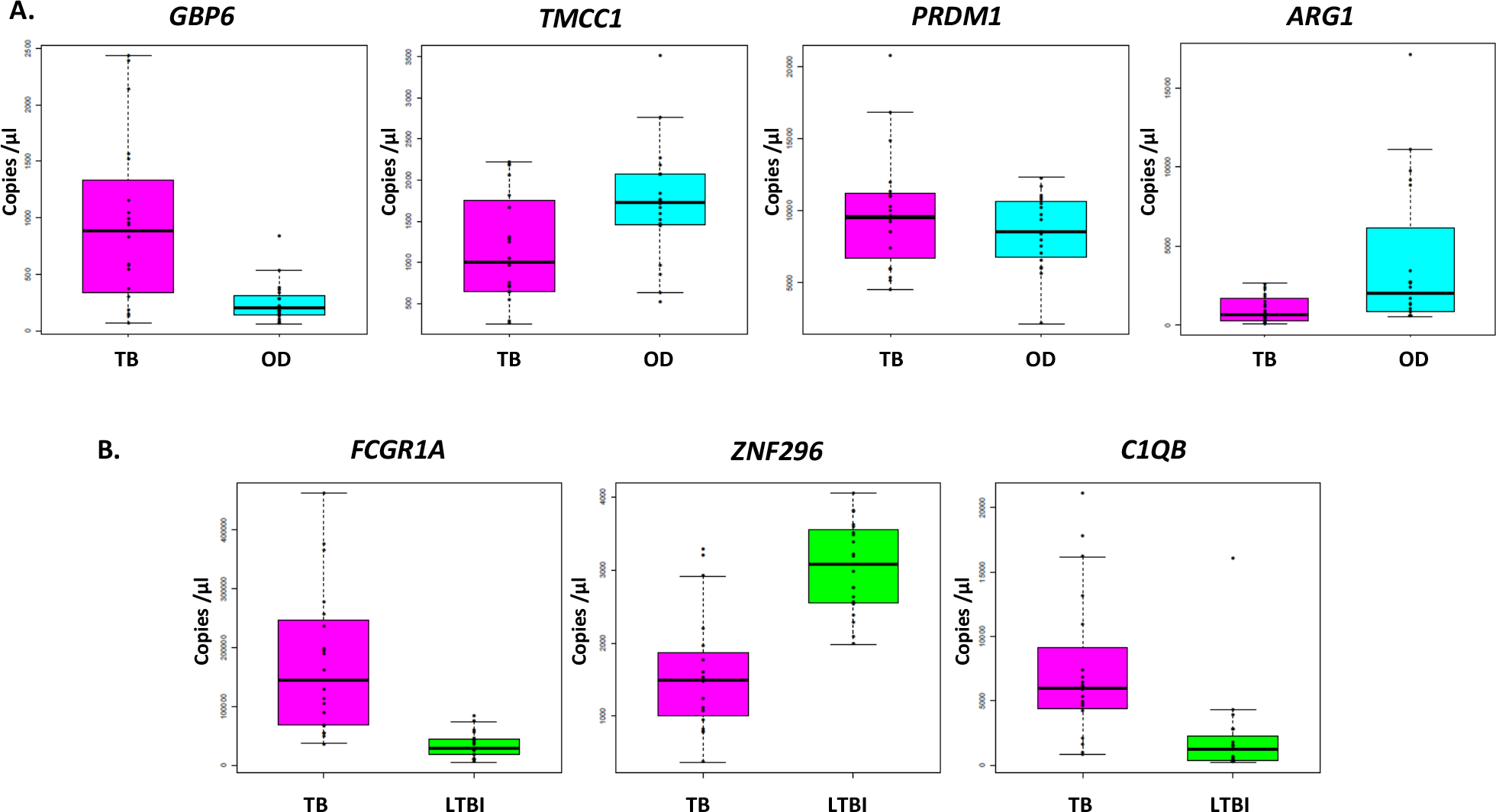
Absolute quantification, expressed as copies per µL for each signature transcript, according to disease group. (A) TB/OD signature transcripts. (B) TB/LTBI signature transcripts. Culture confirmed TB cases are shown in pink (n _TB_ = 20), OD cases in cyan (n _OD_ = 20) and LTBI individuals in green (n _LTBI_ = 20).

### Absolute quantification of genes comprising the three-transcript FS-PLS signature for TB/LTBI

The concentrations (in copies per µL) of each of the transcripts comprising the FS-PLS signature for TB/LTBI in purified RNA from whole blood are shown in Fig 3B. The genes *FCGR1A* and *C1QB* are more abundant in patients with TB compared to LTBI, whereas *ZNF296* is downregulated. All three genes were identified in the original 27 TB/LTBI signature [9]. S2 Fig shows the concentration (in copies per μL) for the samples stratified by HIV status.

### Performance of the four-transcript FS-PLS signature for TB/OD using RT-dPCR analysis disease classification in HIV-infected and HIV-uninfected individuals

The performance of the FS-PLS signature for TB/OD was evaluated by using the DRS to the absolute concentration values that were derived from the RT-dPCR data. Fig 4 (A-D) shows the cross-platform and cross-sample performance of the four gene signature DRS in TB vs OD. In the combined SA/Malawi HIV-infected and -uninfected cohort, the signature had an AUC of 93.8% (CI_95%_: 82.2 – 100), sensitivity 95.0% (CI_95%_: 85.0 – 100) and specificity of 85.0% (CI_95%_: 75.0 – 100) (Fig 4A,4B and Table S5). As observed previously, the accuracy of classification varied with HIV status. The four gene TB/OD signature had an AUC of 91.0% (CI_95%_: 73.3 – 100%) among the HIV-uninfected individuals, and an AUC of 93.0% (CI_95%_: 82.4 – 100%) for the HIV-infected cohort (Fig 4C, 4D).

**Fig 4.**
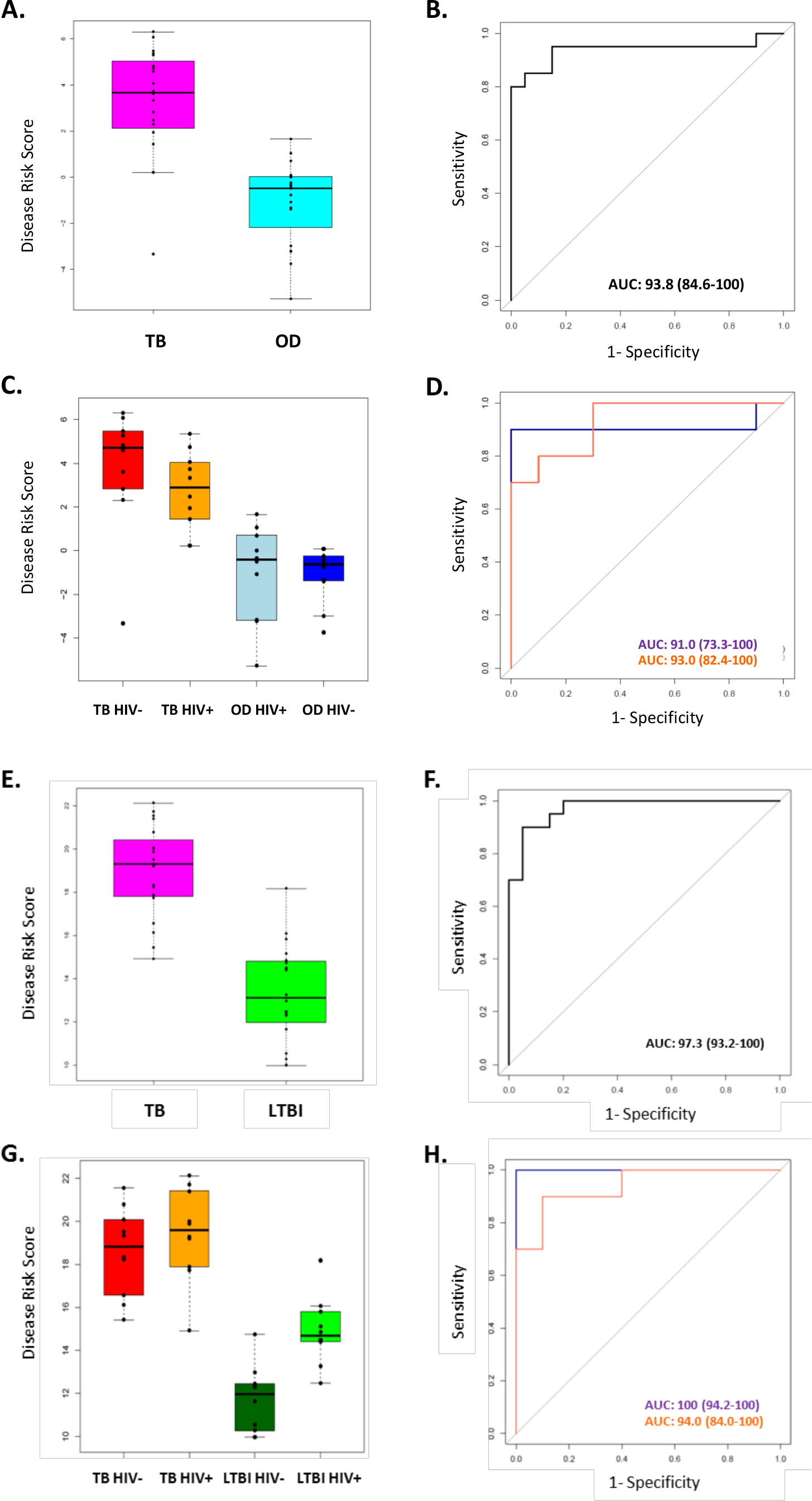
Classification of the SA/Malawi cohorts using the DRS based on the FS-PLS signature RT-dPCR results. (A) Box plots of DRS and (B) ROC curve based on the TB/OD FS-PLS signature applied to the combined TB and OD cohorts (n _TB_ = 20, n _OD_ =20). (C) Box plots of DRS and (D) ROC curve for TB/OD signature, according to HIV infection status (n _TB HIV−_ = 10, n _TB HIV+_ = 10, n _OD HIV−_ = 10, n _OD HIV+_ = 10). (E) Box plots of DRS and (F) ROC curve based on the TB/LTBI FS-PLS signature applied to the combined TB and LTBI cohorts (n _TB_ = 20, n LTBI = 20). (G) Box plots of DRS and (H) ROC curve for TB/LTBI signature, according to HIV infection status (n _TB HIV−_ = 10, n _TB HIV+_ = 10, n _LTBI HIV−_= 10, n _LTBI HIV+_ =10). 95% confidence intervals are shown in brackets.

### Performance of the four-transcript FS-PLS signature for TB/LTBI using dPCR analysis

The performance of the FS-PLS signature for TB/LTBI was evaluated by using the DRS to the absolute log_2_ transformed concentration values that were derived from the RT-dPCR data. Fig 4E-H show the cross-platform and cross-sample performance of the three gene signature DRS in TB vs LTBI. In the combined SA/Malawi HIV-infected and uninfected cohort the signature had an AUC of 97.3% (CI_95%_: 93.3 – 100%), sensitivity of 95.0% (CI_95%_: 85.0 – 100) and specificity of 85.0% (CI_95%_: 75.0 – 100) (Fig 4E and 4F).

As observed previously, the accuracy of classification varied with HIV status. The four gene TB/LTBI signature had an AUC of 100% (CI_95%_: 94.2% – 100%) among the HIV-uninfected individuals and an AUC of 94.0% (CI_95%_: 84.1 – 100%) among HIV-infected cohort (Fig 4G, 4H and S5 Table).

### Correlation of the microarray intensity values and the RT-dPCR concentration values

The expression profiles of the seven genes comprising the two signatures described above were compared between the two platforms, at individual sample level. High correlations were observed between the gene expression profiles generated by the two platforms for most of the genes (Fig 5.). However, differences in expression profiles were also apparent between the two platforms, with a number of samples/genes exhibiting relatively higher expression values in either platform. Pearson correlation and *p* values for all the genes can be found in S6 Table.

**Fig 5.**
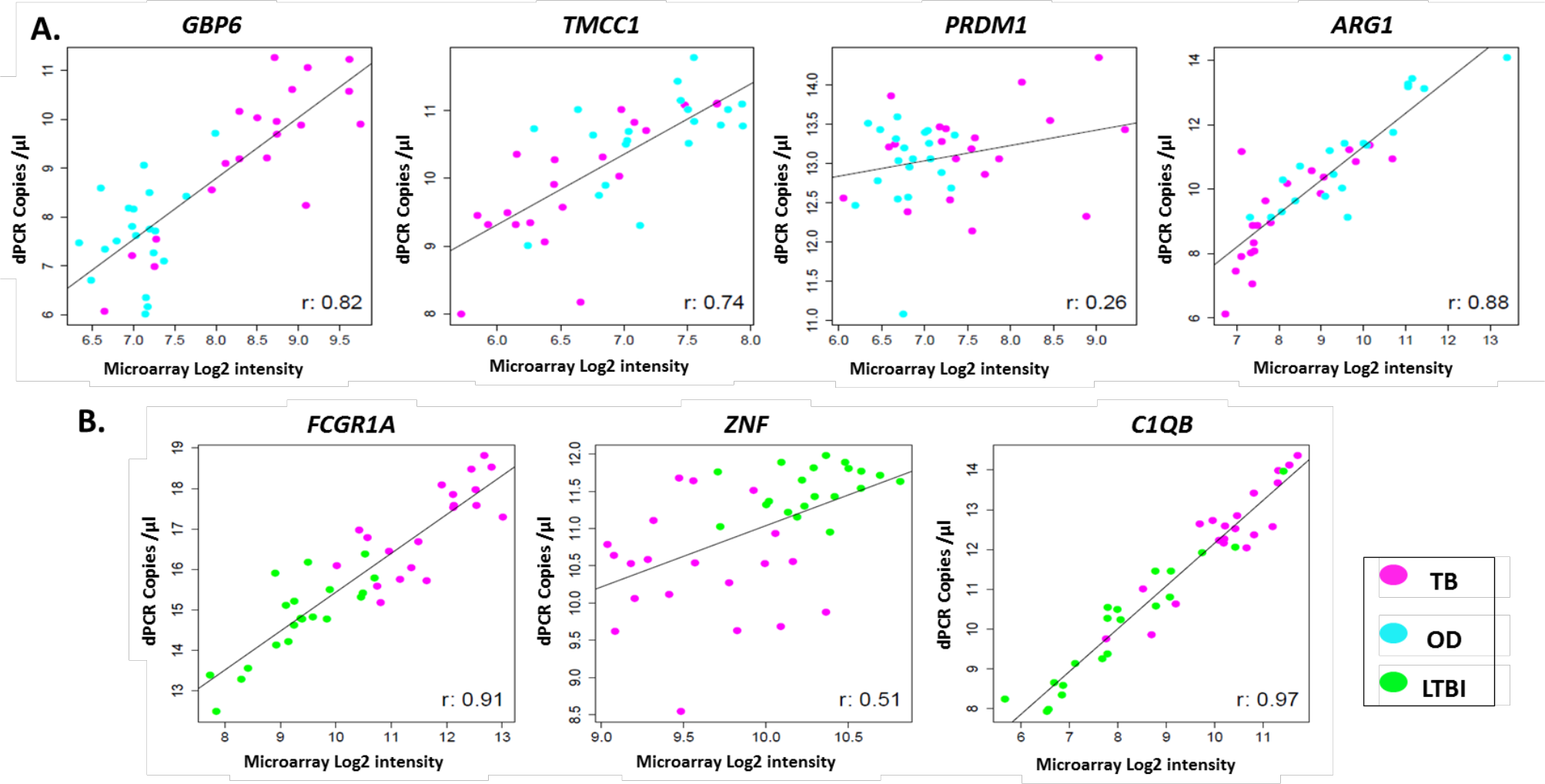
Comparison of expression profiles of common genes and samples between the two platforms. Scatter plots show log_2_ transformed expression values between two platforms per individual. Black line represents line of best fit. for the best-fit line.

## Discussion

In this study, we report a four-gene signature discriminating TB from OD (TB/OD) and a three-gene signature discriminating TB from LTBI (TB/LTBI), which were identified by applying an advanced methodology, FS-PLS, significantly furthering previous landmark work in TB transcriptomics [9] [17] [31] [26]. The performance of the two novel transcriptomic signatures, for TB/OD and TB/LTBI was assessed in the 20% test set and publicly available cohorts. The two signatures were subsequently validated using RT-dPCR and samples from the test cohort, confirming their accuracy of patient classification. We also report estimates for the abundance of each of the individual transcripts in the signatures in purified RNA from whole blood. A weighted regression model was not used in this work, reducing the risk of overfitting and providing more flexibility for application transfer in different detection platforms. This work provides compelling evidence of the robustness and reproducibility of the FS-PLS signatures and the DRS in classifying patients with TB, OD and LTBI and the results presented here support the excellent discriminatory power of both the small gene number TB/OD and TB/LTBI FS-PLS signatures. The point estimates of sensitivity and specificity for our FS-PLS, expressed as DRS and measured by RT-dPCR are within the WHO TPP recommendations [3].

To our knowledge, this study is the first example of the use of RT-dPCR for absolute quantification of transcriptomic signatures in infectious diseases, as anticipated by review articles [32]. Previous studies showed that RT-dPCR has a high accuracy for assessing absolute quantification of RNA and did not show significant inter-assay agreement [22]. However, it should be noted that the efficiency of reverse transcriptase enzymes can be extremely variable and future investigations will be needed to provide further information on absolute abundances of individual RNA transcripts in purified RNA from whole blood. Nevertheless, the concentration values reported in this study provide novel insights that could be of significant use to the diagnostics development research community, providing information regarding the required limits of detection and dynamic range for assays designed to detect signature transcripts. Although high correlation was observed between the gene/sample measurements for the two platforms for most of the genes, the differences reported highlight that a larger number of candidates need to be screened with technology reflective of the point-of-care platform to ensure maximum diagnostic potential.

Clinical applications of dPCR exploit its ability to perform absolute quantification of nucleic acids without the need for rigorous calibration or standardisation between laboratories. RT-dPCR and dPCR have been used to determine copy numbers for a range of pathogens, including the hepatitis B virus, HIV, *Mycobacterium tuberculosis*, *Helicobacter pylori* and *Plasmodium* spp. [23]. While dPCR is more technologically advanced than qPCR, offering absolute rather than relative quantification, the implementation of dPCR in clinical laboratories has been impeded by its relatively low throughput, higher complexity and cost. However, as new instrumentation for dPCR becomes more widely available and simpler to use, it is highly likely that it will play a key role in diagnostic laboratories in the near future [23].

Out of the four transcripts in the TB/OD transcript signature *GBP6* and *PRDM1* are upregulated, and *TMCC1* and *ARG1* are downregulated, in patients with TB compared to OD. *GBP6* appears in the previously published gene expression signature for TB [9], as do other members of the gene family, such as *GBP5* [5] [8] [16] [17] [18], *GBP1* [17] [18] and *GBP2* [18]. These genes are induced by the interferon (IFN) cytokine family [33] and have been shown to be important for cell-autonomous defence against intracellular pathogens [34]. *PRDM1* encodes a DNA-binding protein that acts as a transcriptional repressor of various genes, including IFN-β [35] by binding specifically to the PRDI (positive regulatory domain I element). *PRDM1* has also been shown to regulate the differentiation of B cells into plasma cells that produce antibodies, as well as myeloid cells, such as macrophages and monocytes [36]. Little is known about the function of *TMCC1* in TB pathogenesis, but expression of *ARG1* is induced by toll-like receptor signalling in macrophages [37]. ARG1 plays an important role in the production of nitric oxide (NO), used to kill intracellular pathogens, when nitric oxide synthase-2 (NOS2) is unable to metabolise arginine in hypoxic environments, such as the granuloma [38]. ARG1 is able to produce NO in the absence of oxygen and is therefore critical for the control of intracellular TB [39].

The three gene signature for TB/LTBI reported here consists of two genes that are upregulated (*FCGR1A* and *C1QB*) and one gene that is downregulated (*ZNF296*) in TB compared to LTBI. *FCGR1A* appears in other gene expression signatures for TB [5] [9] [17] [18], and was the most discriminatory gene in a three-gene signature for TB/LTBI [6]. Fc receptors (FcR) play an important role in regulating the immune system and are expressed by a number of innate immune effector cells, particularly monocytes, macrophages, dendritic cells, basophils and mast cells [40]. It has been shown that the monocytic THP-1 cell line upregulates surface expression of Fcγ-RI in response to IFN-γ [41]. *C1QB* encodes a component of the complement 1 (C1Q) complex, part of the complement immune system. Expression of genes encoding components of C1Q have been shown to correlate with the progression of active TB compared to HC and LTBI cohorts [42] and a recent study showed that, in four independent cohorts, components of the C1Q complex are elevated in patients with active TB compared to those with LTBI [43]. *ZNF296* encodes a member of the C2H2 zinc-finger protein family, which contain DNA binding motifs often found in transcription factors. A microarray study identified this gene as upregulated in response to viral infection [44].

This study has certain limitations. The 95% confidence intervals reported for the classification measures for the RT-dPCR validation are relatively wide, due to the small sample size used. 1It is widely accepted that TB diagnosis using transcriptomic signatures offers a number of clear advantages over various sputum-based techniques. However, there are a number of technical challenges of detecting mRNA from whole blood, including sample processing to extract mRNA transcripts that is generally intracellular and inherently less stable than DNA, and that can vary in concentrations by multiple orders of magnitude between samples.

The gene expression signatures for TB/LTBI and TB/OD reported in this study represent extremely promising biomarkers for TB, particularly since they can be measured in whole blood and comprise few analytes. A number of technologies exist that might facilitate their translation into a test, which could include the use of nanomaterials, to quantify mRNA transcripts without an amplification step [45]. A whole blood-based diagnostic test for TB would transform the diagnostic pipeline and enable earlier treatment commencement for patients that would otherwise be missed, and thus prevent onward transmission of the disease, contributing towards paving the way for the end of the TB epidemic by 2030, Goal 3.3 of the Sustainable Development Goals, as set out by the United Nations [46].

## Supporting information

MIQEchecklist

STARDchecklist

Raw dPCR data

## Acknowledgements

The study was funded by an EU Action for Diseases of Poverty program grant (Sante/2006/105-061) and made use of infrastructure and staff at the Wellcome Trust–supported programs in Karonga and University of Cape Town and the Imperial College Centre for Clinical Tropical Medicine. The Karonga Prevention Study is supported by the Wellcome Trust, UK (079828/079827); RJW and AJB were supported by the Wellcome Trust, UK (104803 and 203135). RJW is also supported by the Francis Crick Institute which receives its core funding from Cancer Research UK (FC00110218), the UK Medical Research Council (FC00110218), and The Wellcome Trust (FC00110218). The funders had no role in study design, data collection and analysis, decision to publish, or preparation of the manuscript. ML and MK receive support from the Imperial College BRC. MK also acknowledges support from the Wellcome Trust (Sir Henry Wellcome Fellowship grant 206508/Z/17/Z). HDG acknowledges support from EPSRC through the Interdisciplinary Research Centre (IRC) “Early-Warning Sensing Systems for Infectious Diseases” (EP/K031953/1).

