## Supplementary material for "Identification of reduced host transcriptomic signatures for tuberculosis and digital PCR-based validation and quantification": Suplementary

### Supporting Information Appendix

**ILULU Consortium members:**

**Institute of Infectious Diseases and Molecular Medicine, University of Cape Town** Nonzwakazi Bangani, Lizl Bashe, Melina Carr, Hannah P. Gideon, Rene Goliath, Yekiwe Hlombe, Vanessa January, Bekekile Kwaza, Suzaan Marais, Marc Mendelson, Tolu Oni, Fadheela Patel, Ronnett Seldon, Relebohile Tsekela, Katalin A. Wilkinson, Robert J. Wilkinson, Kathryn Wood; **London School of Hygiene and Tropical Medicine/Karonga Prevention Study** Lyn Ambrose, Amelia C. Crampin, Hazel M. Dockrell, Neil French, Lumbani Munthali, Bagrey Ngwira, Amos Phiri, Femia Zgambo; **Red Cross War Memorial Children's Hospital, University of Cape Town** Margaret Cooper, Brian Eley, Mabel Gcuwa, Spasina King, Glynis Kossew, Karen McCabe, Wonita Petersen, Sandra Pienaar, Vashini Pillay; **Liverpool School of Tropical Medicine/Malawi-Liverpool-Wellcome Trust Clinical Research Programme**, **University of Malawi College of Medicine** Benjamin Allubha, George Chagaluka, Angeziwa Chunga, Janet Dube, Robert S. Heyderman, Annie Joabe, Martha Kalemba, Anne Kerr, Monica Matola, Rachel Mlotha, Agnes Mwale, David Mzinza; **Brighton and Sussex Medical School, University of Sussex** Suzanne T. Anderson, Gillian Baker, Claire M. Banwell, Terry Bishop, Natalie Chaplin, Julian Golland, Florian Kern, Susan Poore, Jayne Wellington; **Imperial College London** Lachlan J. Coin, Harieta Eleftherohorinou, Shea Hamilton, Harriet D. Gliddon, Dominic Habgood-Coote, Myrsini Kaforou, Paul R. Langford, Michael Levin, Stephanie Menikou, Victoria J. Wright.

#### Supplementary Methods

##### *Recruitment of patients*

**Definite TB case:** a participant with a clinical condition consistent with tuberculosis and microbiological confirmation with evidence from at least two specimens confirming the presence of AFB with at least one specimen confirmed on culture as MTB complex.

**Latent TB infected (LTBI) case:** a participant who is clinically assessed as healthy and not suffering from a clinical syndrome in which tuberculosis is likely. The individuals will have a tuberculin skin test (TST) size of 10mm or more if HIV negative, or 5mm or more if HIV positive and a positive Interferon Gamma Release Assay (IGRA) and negative sputum culture. Sputums were only collected in Malawi if cough was productive, when at least two samples would be collected. LTBI criteria were later relaxed to allow a positive TST and/or a positive IGRA to facilitate recruitment in Malawi. This change was made prior to any gene expression measurements.

**Other disease (OD) case:** A participant with a disease syndrome that on presentation includes tuberculosis in the differential diagnosis, but following clinical management will have tuberculosis excluded and a firm alternative diagnosis established.

##### *RNA sample extraction and processing*

Whole blood (2.5ml) was collected into PAXgene blood RNA tubes (PreAnalytiX, Germany), incubated for 2 hours, frozen at -20$^{o}$C within 3 hours of collection, and then stored at -80$^{o}$C. RNA was extracted using PAXgene blood RNA kits (PreAnalytiX, Germany) according to the manufacturer's instructions at one site (Cape Town) to minimize any sample handling bias. The integrity and yield of the total RNA was assessed using an Agilent 2100 Bioanalyser and a NanoDrop 1000 spectrophotometer respectively. The mean RIN for the purified RNA samples was 7.93 ± 0.78, with a slightly higher RIN measured for samples from South Africa (8.45, SD 0.37) compared to those from Malawi (7.61, SD 0.79). The average rRNA ratio (28S/18S) was 1.68 ± 0.46.

##### *dPCR FAM fluorescence threshold setting*

In dPCR, FAM call thresholds decide which dPCR reactions are classified as negative (below the threshold) and which are classified as positive (above the threshold). Although it is preferable for the FAM call thresholds to be fixed for a given TaqMan assay, this was not possible for the *GBP6*, *TMCC1* and *ZNF296* probes, due to variation in FAM fluorescence intensities between and within dPCR runs. The x- and y-axes of the FAM fluorescence plots on the QuantStudio 3D AnalysisSuite Cloud Software Version 3.0.3 (Thermo Fisher Scientific) are not fixed, and the axes are automatically adjusted for each chip. Differences in the optimal placing of FAM call thresholds can also be due to sub-optimal amplification efficiencies of the TaqMan assays. For example, S1 Fig shows the results when the TaqMan assay for *GBP6* was used in a dPCR with three different samples. S1A Fig shows the results of a no template control. The automatic threshold of 6063 identified all partitions as having negative reactions (in yellow) except for one (in blue), which showed a fluorescence intensity level above the threshold. The thresholds for other samples varied according to how the positive and negative reaction populations separated. The threshold for sample 20311 was automatically set at 7291 by the software, but 7417 was subsequently manually set because it achieved improved separation of the populations (S1B Fig). The threshold for sample 20665 was automatically set at 5527 (S1C Fig).

To ensure that there was no systematic bias in assigning higher or lower FAM call thresholds to either the TB, LTBI or OD cohorts, it was verified that the thresholds did not vary by disease grouping. The mean and range FAM call threshold for each assay were compared between each group and can be reviewed in S4 Table. Similarly, the lambda value, volume and dilution of sample used, and FAM call thresholds for each sample and probe combination are shown in S1 Fig in accordance with the MIQE guidelines [1].

***RT-dPCR data analysis***

Two reference genes, *β-actin* and 18s rRNA were included in this work in order to provide additional quality control, but were not used for normalisation or the derivation of the disease risk score (DRS). The *β-actin* gene has been used as an endogenous control for normalisation in TB studies [2] [3]. The levels of the 18S transcript vary according to HIV infection status and TB status, with a higher concentration observed in HIV-uninfected TB patients. The levels of the β-actin transcript also varied across the disease groups, with lower levels observed in HIV-infected TB patients (S1 Fig). For this reason, $\log_2$ concentration values for each of the signature transcripts were used to calculate the DRS, without normalisation using a reference gene.

The Poisson statistics used to analyse dPCR data assumes that every molecule of DNA has an equal chance of entering any one of the partitions on a dPCR chip. The mean copies per partition, known as λ, can then be calculated. In order to achieve the highest precision, it is advised that the concentration of target that is added to each individual reaction is between 140 and 3,070 copies/µL, which correspond to λ values of 0.1 and 2.3 and Poisson-corrected values of 2,000 to 45,000 copies/chip. In some cases, where the concentration of a particular target was extremely low, this was not possible and the λ value fell below 0.2. In these cases, the maximum amount of sample was used (5 µL). In these cases, the results may be less precise because of the way the Poisson statistics are applied to the data. However, it has been shown that the reproducibility and accuracy of the concentration values where the λ value is less than 0.2 is generally very good. However, where the λ value exceeds 3, systematic bias is observed (unpublished data), and so this was avoided. The median λ values for each TaqMan assay and each disease cohort are summarised in S6B Table.

***Absolute quantification of RNA concentration***

For each chip, the number of negative partitions (*z*) and the total number of partitions (*n*) was taken from the analysis of the QuantStudio 3D AnalysisSuite Cloud Software Version 3.0.3. From this, the number of positive partitions (*k*) could be calculated using the simple formula:

$$k = n - z$$

The λ values could then be calculated by:

$$\lambda = -\ln (1 - z/n)$$

The concentration (C) was then calculated using the following formula:

$$C = \frac{\lambda}{V}$$

Where *V* is the partition volume, which for the QuantStudio 3D Digital PCR 20K Chips, was 809 pL. This gave a concentration value in copies/µL, which was then corrected for depending on the volume and dilution of sample used, so these values indicate the concentration of each gene of interest (GOI) in the RT mix. In order to identify the concentration of each GOI in purified RNA from whole blood, the dilution and volume of sample used for dPCR was corrected for. Then, the amount of purified RNA sample used for the RT reaction was corrected for, so the final value gave an indication of the GOI concentration in purified RNA from whole blood. To transfer from copies/µL to moles/L (molarity, M), we multiply by 1,000,000 and divide by Avogadro's constant ($6.022 \times 10^{23}$ $moles^{-1}$). Despite these concentrations being calculated using the amplicon molecular weight, they also hold true for the mRNA species from which the amplicon was made.

$$\text{Conc. in whole blood RNA (copies/µL)} = \frac{\text{Conc. (copies/µL) in dPCR} \times 20\ \text{µL (total dPCR vol.)}}{\text{Vol. of whole blood RNA used for dPCR (µL)}}$$

$$\text{Conc. in whole blood RNA (mole/L)} = \frac{\text{Conc. in whole blood RNA (copies/L)}}{6.02 \times 10^{23}\ \text{(copies/mole) (Avogadro's number)}}$$

#### Supplementary Statistical Methods

##### *Microarray analysis using FS-PLS*

Mean raw intensity values for each probe were corrected for local background intensities and a robust spline normalisation [4] (combining quantile normalisation and spline interpolation) was applied to each array. Expression values were transformed to a logarithmic scale (base 2), and for each probe. Differential expression between patient groups was identified by fitting a linear model to each transcript using LIMMA [5]. P-values were adjusted using the method of Benjamini and Hochberg [6]. Transcripts with log FC >0.5 were taken forward to variable selection with FS-PLS [7] [8]. This threshold was chosen in order to ensure that differential expression for selected variables could be distinguished using the resolution of RT-PCR.

##### *Disease risk score (DRS) calculation*

For each individual, we calculated the DRS using the minimal transcript selected sets for TB vs. LTBI and TB vs. OD. The $\log_2$ expression value of the transcripts identified by FS-PLS were combined into a DRS without weights to only denote the direction of expression. The score is based on subtracting the summed intensities of the down-regulated transcripts from the summed intensities of the up-regulated transcripts, as described previously [9].

##### *Analysis of validation datasets*

For validation of the performance of the DRS based on the TB/OD four transcript and TB/LTBI 3 transcript signature, we used the whole blood expression datasets of Berry et al. [10] and Bloom et al. [11] generated using Illumina HT12 Beadarrays (accession series GSE19491, GSE42834). The cohorts comprised HIV-uninfected individuals; including TB, LTBI, Healthy Controls and other diseases including pneumonia, lung cancer, Still's disease, Systemic Lupus Erythematosus, *Staphylococcus*, and *Streptococcus*. For the evaluation of the performance of our TB/LTBI signature in the Berry et al. [10] UK training and test datasets, we combined them using Combat, to account for the observed batch effect [12].

#### Supplementary Results

***S1 Table. Statistical power calculations in order to specify minimum sample size for dPCR were calculated according to the required sensitivity of dPCR results in classifying patients.***

| | | dPCR sensitivity | | | |
|---|---|---|---|---|---|
| | | **0.7** | **0.75** | **0.8** | **0.85** |
| **Level of statistical significance** | **0.05** | n=50 | n=31 | n=21 | n=14 |
| | **0.1** | n=38 | n=24 | n=15 | n=11 |

***S2 Table. The clinical diagnoses of the OD cohort.***

| **HIV infection status** | **HIV infected** | | **HIV uninfected** | | **Total** |
|---|---|---|---|---|---|
| **Location** | **SA** | **MLW** | **SA** | **MLW** | |
| **Pneumonia/LRTI/PJP** | 2 | 1 | 1* | 1 | 5 |
| **Malignancy and other neoplasia other than Kaposi's sarcoma** | | | 4* | | 4 |
| **Pelvic inflammatory disease/UTI** | 1 | | 2 | 3 | 6 |
| **Bacterial, viral meningitis, or meningitis of uncertain origin** | 1 | 2 | | | 3 |
| **Kaposi's sarcoma** | 1 | | | | 1 |
| **Gastric ulcer or gastritis** | 1 | | | | 1 |
| **Total** | 6 | 4[‡] | 6 | 4 | 20 |

***S3 Table. TaqMan assays used for dPCR. All were inventoried and available from Applied Biosystems.***

| Gene | TaqMan probe used for dPCR | RefSeq or GenBank ID | Exon boundary | Amplicon length |
|---|---|---|---|---|
| *GBP6* | Hs01584201_m1 | NM_001320257.1 | 8-9 | 111 |
| | | NM_198460.2 | 10-11 | 111 |
| | | XM_011540835.2 | 10-11 | 111 |
| | | AK131329.1 | 5-6 | 111 |
| | | AK131356.1 | 10-11 | 111 |
| | | AK290303.1 | 10-11 | 111 |
| | | AK299408.1 | 9-10 | 111 |
| | | AK299466.1 | 8-9 | 111 |
| | | AK303987.1 | 10-11 | 111 |
| | | AK315834.1 | 10-11 | 111 |
| | | BC131713.1 | 10-11 | 111 |
| | | BX537949.1 | 10-11 | 111 |
| | | BX647907.1 | 10-11 | 111 |
| *TMCC1* | Hs01037666_s1 | NM_001017395.3 | 6 | 129 |
| | | NM_001128224.2 | 6 | 129 |
| | | NR_033361.1 | 4 | 129 |
| | | XM_006713542.3 | 7 | 129 |
| | | XM_006713543.3 | 7 | 129 |
| | | XM_006713544.1 | 7 | 129 |
| | | XM_006713550.3 | 7 | 129 |
| | | XM_006713552.3 | 4 | 129 |
| | | XM_006713553.3 | 4 | 129 |
| | | XM_011512572.2 | 8 | 129 |
| | | XM_011512573.2 | 7 | 129 |
| | | XM_011512576.2 | 7 | 129 |
| | | XM_011512579.2 | 6 | 129 |
| | | XM_011512580.2 | 9 | 129 |
| | | XM_011512581.2 | 8 | 129 |
| | | XM_011512582.1 | 4 | 129 |
| | | XM_011512583.2 | 6 | 129 |
| | | XM_011512584.2 | 8 | 129 |
| | | XM_017005931.1 | 7 | 129 |
| | | XM_017005932.1 | 8 | 129 |
| | | XM_017005933.1 | 6 | 129 |
| | | XM_017005934.1 | 10 | 129 |
| | | XM_017005935.1 | 11 | 129 |
| | | XM_017005936.1 | 8 | 129 |
| | | XM_017005937.1 | 8 | 129 |
| | | XM_017005938.1 | 10 | 129 |
| | | XM_017005939.1 | 9 | 129 |
| | | XM_017005940.1 | 6 | 129 |
| | | XM_017005941.1 | 6 | 129 |
| | | CR749206.1 | 4 | 129 |
| *PRDM1* | Hs00153357_m1 | NM_001198.3 | 4-5 | 65 |
| | | NM_182907.2 | 2-3 | 65 |
| | | XM_006715550.3 | 4-5 | 65 |
| | | XM_011536062.2 | 4-5 | 65 |
| | | XM_011536063.2 | 4-5 | 65 |
| | | XM_011536064.2 | 2-3 | 65 |
| | | XM_017011187.1 | 4-5 | 65 |
| | | AF084199.1 | 4-5 | 65 |
| | | AK289556.1 | 4-5 | 65 |
| | | AK301341.1 | 2-3 | 65 |
| | | AL832963.1 | 4-5 | 65 |
| | | AY198414.1 | 4-5 | 65 |
| | | AY198415.1 | 2-3 | 65 |
| | | BC103832.1 | 2-3 | 65 |
| | | BC103833.1 | 2-3 | 65 |
| | | BC103834.1 | 2-3 | 65 |
| | | BC103835.1 | 2-3 | 65 |
| *ARG1* | Hs00968979_m1 | NM_000045.3 | 4-5 | 78 |
| | | NM_001244438.1 | 4-5 | 78 |
| | | AY074488.1 | 4-5 | 78 |

| | | | | |
|---|---|---|---|---|
| | | BC005321.1 | 4-5 | 78 |
| | | BC020653.1 | 4-5 | 78 |
| | | BG217880.1 | 4-5 | 78 |
| | | BG542163.1 | | 78 |
| | | BT006741.1 | 4-5 | 78 |
| | | M14502.1 | 4-5 | 78 |
| *FCGR1A* | Hs02340031_m1 | NM_000566.3 | 5-6 | 151 |
| | | XM_005244957.3 | 5-6 | 151 |
| | | XM_005244958.4 | 3-4 | 151 |
| | | AK291451.1 | 5-6 | 151 |
| | | AK291502.1 | 5-6 | 151 |
| | | BC032634.1 | 5-6 | 151 |
| | | BC152383.1 | 5-6 | 151 |
| | | DQ786309.1 | 3-4 | 151 |
| | | L03418.1 | 5-6 | 151 |
| | | X14355.1 | 5-6 | 151 |
| | | X14356.1 | 5-6 | 151 |
| *ZNF296* | Hs00377132_m1 | NM_145288.1 | 2-3 | 74 |
| | | AF447583.1 | 3-4 | 74 |
| | | AY040679.1 | 1-2 | 74 |
| | | BC019352.1 | 2-3 | 74 |
| *C1QB* | Hs00608019_m1 | NM_000491.3 | 2-3 | 78 |
| | | XM_011542059.2 | 3-4 | 78 |
| | | BC008983.1 | 2-3 | 78 |
| | | BQ711602.1 | 3-4 | 78 |
| | | X03084.1 | 1-2 | 78 |
| 18S | Hs99999901_s1 | X03205.1 | 1 | 187 |
| *β-actin* | Hs99999903_m1 | NM_001101.3 | 1 | 171 |
| | | AK025375.1 | 1 | 171 |
| | | AK058019.1 | 1 | 171 |
| | | AK130062.1 | 1 | 171 |
| | | AK130157.1 | 1 | 171 |
| | | AK222925.1 | 1 | 171 |
| | | AK223032.1 | 1 | 171 |
| | | AK223055.1 | 1 | 171 |
| | | AK225414.1 | 1 | 171 |
| | | AK301372.1 | 1 | 171 |
| | | AK304552.1 | 1 | 171 |
| | | AK309997.1 | 1 | 171 |
| | | BC001301.1 | 1 | 171 |
| | | BC002409.2 | 1 | 171 |
| | | BC013380.2 | 1 | 171 |
| | | BC013835.1 | 1 | 171 |
| | | BC014861.1 | 1 | 171 |
| | | BC016045.1 | 1 | 171 |
| | | X63432.1 | 1 | 171 |

***S4 Table. Digital PCR FAM call thresholds, λ values, volumes and dilutions of sample used.***

The median and range of (a) the FAM call thresholds, (b) λ values and (c) volume and dilution of sample used for each TaqMan assay was compared for each cohort.

(a)

| **Assay** | **FAM call threshold median (range)** | | |
|---|---|---|---|
| | **TB Cohort** | **LTBI Cohort** | **OD Cohort** |
| **FCGR1A** | 3000 (3000) | 3000 (3000) | / |
| **ZNF296** | 2025 (1875-2025) | 2025 (1900-2025) | / |
| **C1QB** | 4250 (4250) | 4250 (4250) | / |
| **GBP6** | 6623 (5553-7586) | / | 6623 (5097-7892) |
| **TMCC1** | 3200 (3200) | / | 3200 (3200) |
| **PRDM1** | 4182 (3140-4933) | / | 4277 (3200-5834) |
| **ARG1** | 4000 (4000) | / | 4000 (4000) |
| **18s** | 4000 (4000) | 4000 (4000) | 4000 (4000) |
| **B-actin** | 2600 (2600) | 2600 (2600) | 2600 (2600) |

(b)

| **Assay** | **λ median (range)** | | |
|---|---|---|---|
| | **TB Cohort** | **LTBI Cohort** | **OD Cohort** |
| **FCGR1A** | 0.41 (0.13-2.33) | 1.07 (0.20-2.82) | / |
| **ZNF296** | 0.25 (0.10-0.53) | 0.42 (0.27-0.97) | / |
| **C1QB** | 0.75 (0.08-2.85) | 0.22 (0.07-2.17) | / |
| **GBP6** | 0.24 (0.02-0.66) | / | 0.05 (0.02-0.23) |
| **TMCC1** | 0.27 (0.07-0.60) | / | 0.47 (0.14-0.95) |
| **PRDM1** | 0.36 (0.08-2.99) | / | 2.27 (0.06-2.97) |
| **ARG1** | 0.17 (0.02-0.71) | / | 0.54 (0.07-2.99) |
| **18s** | 0.62 (0.31-1.34) | 0.47 (0.16-1.30) | 0.64 (0.19-1.02) |
| **B-actin** | 0.14 (0.06-0.25) | 0.13 (0.01-0.21) | 0.17 (0.01-0.33) |

**(c)**

| **Assay** | **Volume used/μL (dilutions)** | | |
|---|---|---|---|
| | **TB Cohort** | **LTBI Cohort** | **OD Cohort** |
| **FCGR1A** | 2.5 (neat, 1/10 or 1/20) | 2.5 (neat or 1/10) | / |
| **ZNF296** | 2.5 or 5 (neat) | 2.5 or 5 (neat) | / |
| **C1QB** | 2.5 or 5 (neat or 1.20) | 2.5 or 5 (neat) | / |
| **GBP6** | 5 (neat) | / | 5 (neat) |
| **TMCC1** | 5 (neat) | / | 5 (neat) |
| **PRDM1** | 5 (neat, 1/10 or 1/20) | / | 5 (neat, 1/10 or 1/20) |
| **ARG1** | 5 (neat) | / | 5 (neat) |
| **18s** | 1 (1/50,000) | 1 (1/50,000) | 1 (1/50,000) |
| **B-actin** | 1 (1/10,000) | 1 (1/10,000) | 1 (1/10,000) |

***S5 Table. 2x2 Tables for diagnostic accuracy of patient classification using the FS-PLS signatures on the (A) Microarray test set and (B) RT-dPCR data.***

**A**

| **TB/OD** | | Reference | |
|---|---|---|---|
| | | 42 | 34 |
| | | + | - |
| Index | + | 0 | 34 |
| | - | 42 | 0 |

| **TB/LTBI** | | Reference | |
|---|---|---|---|
| | | 37 | 39 |
| | | + | - |
| Index | + | 0 | 39 |
| | - | 37 | 0 |

**B**

| **TB/OD** | | Reference | |
|---|---|---|---|
| | | 20 | 20 |
| | | + | - |
| Index | + | 0 | 20 |
| | - | 20 | 0 |

| **TB/LTBI** | | Reference | |
|---|---|---|---|
| | | 20 | 20 |
| | | + | - |
| Index | + | 0 | 20 |
| | - | 20 | 0 |

***S6 Table. Correlation between microarray and RT-dPCR results: Pearson correlation and p values.***

| | **Pearson's correlation coefficient** | **95% CI** | **p-value** |
|---|---|---|---|
| ***FCGR1A*** | 0.906 | 0.828 - 0.950 | 9.02E-16 |
| ***ZNF296*** | 0.511 | 0.237 - 0.710 | 0.0007551 |
| ***C1QB*** | 0.967 | 0.938 - 0.982 | < 2.2e-16 |
| ***GBP6*** | 0.818 | 0.680 - 0.900 | 1.11E-10 |
| ***TMCC1*** | 0.741 | 0.558 - 0.855 | 4.56E-08 |
| ***PRDM1*** | 0.256 | -0.060 - 0.526 | 0.111 |
| ***ARG1*** | 0.882 | 0.787 - 0.936 | 5.23E-14 |

***Figure S1. QuantStudio dPCR data analysis.***

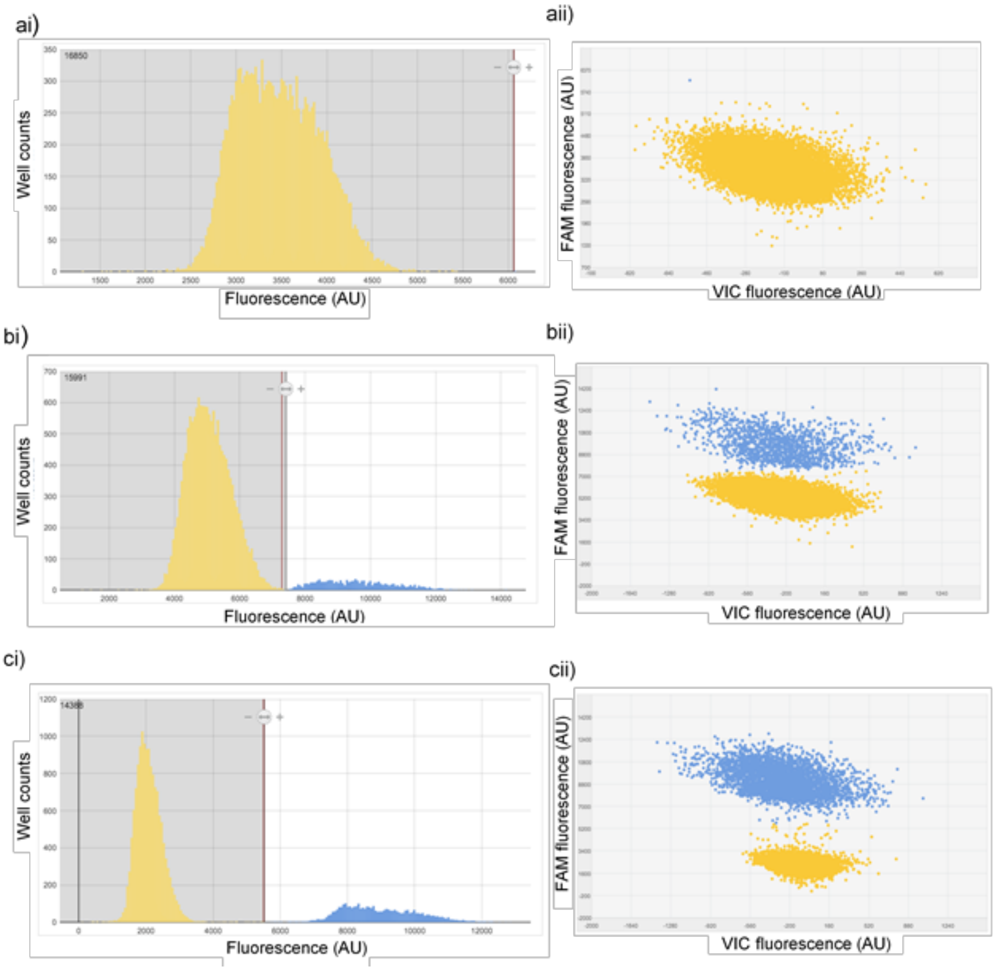

**Figure S1.** QuantStudio dPCR data analysis. The results of three different samples are shown when the GBP6 TaqMan assay was used for dPCR. Positive reactions are depicted in blue and negative reactions in yellow. The threshold is important for accurately separating the two populations of negative and positive reactions. (ai) Histogram view and (aii) Scatter plot view for the no template control. Call threshold was automatically set at 6063. (bi) Histogram view and (bii) Scatter plot view for sample 20311 (SA, TB, HIV+). FAM call threshold was manually set at 7,417. (ci) Histogram view and (bii) Scatter plot view for sample 20665 (SA, OD, HIV+). FAM call threshold was automatically set at 5,527. Y-axes in (aii), (bii) and (cii) range 700 to 7,000 arbitrary values (AU), -2,000 to 16,000 and -2,000 to 16,000 AU respectively. X-axes in (aii), (bii) and (cii) range from -1,000 to 800, -2,000 to 1,600 and -2,000 to 1,600 AU respectively. Scales are set automatically by the software and depend on the range of fluorescence values read on a particular chip.

***Figure S2. Absolute quantification, expressed as copies per µL for each signature transcript, according to disease group stratified by HIV status.***

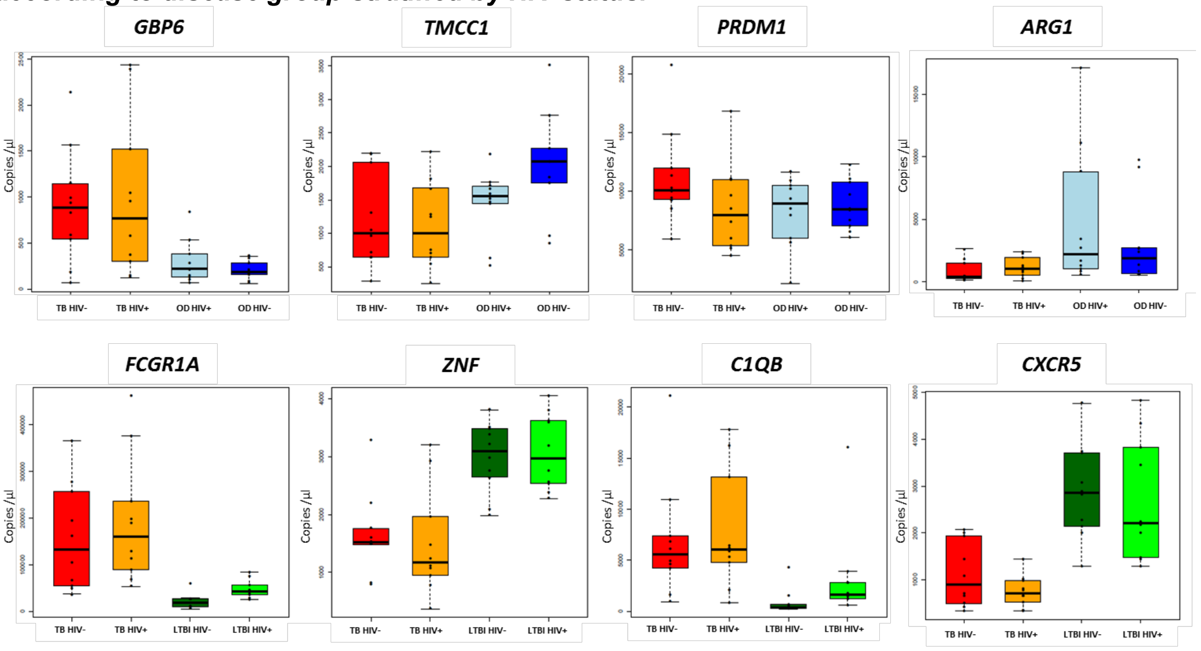

**Figure S2.** Absolute quantification, expressed as copies per µL for each signature transcript, according to disease group stratified by HIV status. Top: TB/OD signature transcripts. Bottom: TB/LTBI signature transcripts. Culture confirmed TB cases HIV uninfected are shown in red ($n_{TB\ HIV-}$ = 10) and HIV infected in orange ($n_{TB\ HIV+}$ = 10), OD cases HIV uninfected in blue ($n_{OD\ HIV-}$ = 10) and HIV infected ($n_{OD\ HIV+}$ = 10) in dark blue, LTBI individuals HIV uninfected in dark green ($n_{LTBI\ HIV-}$ = 10) and HIV infected ($n_{LTBI\ HIV+}$ = 10) in light green.

***Figure S3. Workflow from the original blood sample to the cDNA used for dPCR.***

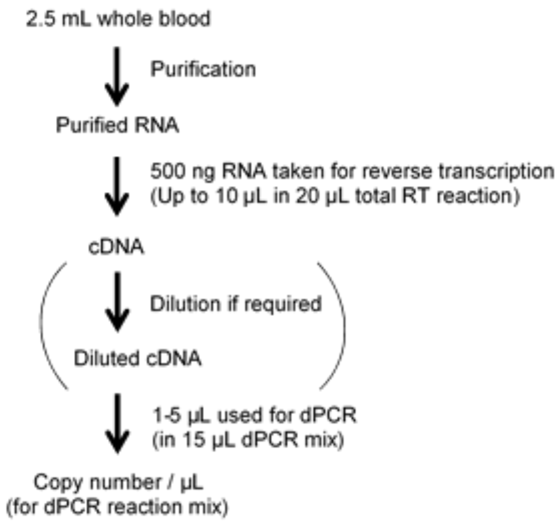

**Figure S3.** Workflow from the original blood sample to the cDNA used for dPCR. After blood samples were taken in a PAXgene tubes, RNA was purified from the sample, followed by reverse transcription to cDNA, which was then used for dPCR.
