## Supplementary material for "Identification of reduced host transcriptomic signatures for tuberculosis and digital PCR-based validation and quantification": MIQEchecklist

| **dPCR MIQE Checklist.**  All essential information (E) must be submitted with the manuscript. Desirable information (D) should be submitted if possible.  When single dPCR experiments are performed, the variation due to counting error alone should be calculated from the Binomial (or suitable equivalent) distribution. | | |
| --- | --- | --- |
| **ITEM TO CHECK** | **IMPORTANCE** | **CHECKLIST** |
| **EXPERIMENTAL DESIGN** |  |  |
| Definition of experimental and control groups | **E** | ✓ page 9 |
| Number within each group | **E** | ✓ page 12 |
| Assay carried out by core lab or investigator's lab? | D |  |
| Power analysis | D | ✓ page 8, 9 |
| **SAMPLE** |  |  |
| Description | **E** | ✓ page 8, 9, 10, 11 |
| Volume/mass of sample processed | D | ✓ page 9 |
| Microdissection or macrodissection | **E** | NA |
| Processing procedure | **E** | NA |
| If frozen - how and how quickly? | **E** | NA |
| If fixed - with what, how quickly? | **E** | NA |
| Sample storage conditions and duration (especially for FFPE samples) | **E** | ✓ page 9, 10 |
| **NUCLEIC ACID EXTRACTION** |  |  |
| Nucleic acid quantification | **E** | ✓ page 9, 10 |
| DNA or RNA quantification | **E** | ✓ page 9, 10 |
| Quality/Integrity, method/instrument, e.g. RNA integrity | **E** | ✓ page 9 |
| Template structural information | **E** | NA |
| Template modification (digestion, sonication, preamplification etc) | **E** | ✓ page 10 |
| Template treatment | **E** | ✓ page 10 |
| Inhibition dilutions or spike | **E** | ✓ page 10 |
| DNA contamination assessment of RNA samples | **E** | NA |
| Details of DNase treatment where performed | **E** | NA |
| Manufacturer of reagents used and catalogue number | D | ✓ page 9 |
| Storage conditions (Nucleic acid): temperature, concentration, duration, buffer) | **E** | ✓ page 9 |
| **REVERSE TRANSCRIPTION (if necessary)** |  |  |
| cDNA priming method and concentration | **E** | ✓ page 10 |
| One or two-step protocol | **E** | ✓ page 10 |
| Amount of RNA used per reaction | **E** | ✓ page 10 |
| Detailed reaction components and conditions | **E** | ✓ page 10 |
| RT efficiency | D |  |
| Estimated copies measured with and without addition of RT | D |  |
| Manufacturer of reagents and catalogue numbers | D | ✓ page 10 |
| Reaction volume | D | ✓ page 10 |
| Storage conditions of cDNA | D | ✓ page 10 |
| **dPCR TARGET INFORMATION** |  |  |
| Sequence accession number | **E** | ✓ S3 Table |
| Location of amplicon | D |  |
| Amplicon length | **E** | ✓ S3 Table |
| *In silico* specificity screen (BLAST, etc) | **E** | NA |
| Pseudogenes, retropseudogenes or other homologs? | D |  |
| Sequence alignment | D |  |
| Secondary structure analysis of amplicon and GC content | D |  |
| Location of each primer by exon or intron (if applicable) | **E** | NA |
| What splice variants are targeted? | **E** | ✓ S3 Table |
| **dPCR OLIGONUCLEOTIDES** |  |  |
| Primer sequences | **E** | ✓ page 10, S3 Table |
| RTPrimerDB Identification Number | D |  |
| Probe sequences | D |  |
| Location and identity of any modifications | **E** | NA |
| Manufacturer of oligonucleotides | D | ✓ page 10 |
| Purification method | D |  |
| **dPCR PROTOCOL** |  |  |
| Complete reaction conditions | **E** | ✓ page 10 |
| Reaction volume and amount of cDNA/DNA | **E** | ✓ page 10 |
| Primer, (probe), Mg++ and dNTP concentrations | **E** | ✓ page 10 |
| Polymerase identity and concentration | **E** | ✓ page 10 |
| Buffer/kit identity and manufacturer | **E** | ✓ page 10 |
| Exact chemical constitution of the buffer | D |  |
| Additives (SYBR Green I, DMSO, etc.) | **E** | ✓ page 10 |
| Plates/tubes catalogue number and manufacturer | D | ✓ page 10 |
| Complete thermocycling parameters | **E** | ✓ page 10 |
| Reaction setup (manual/robotic) | D | ✓ page 10 |
| Gravimetric or volumetric dilutions (manual/robotic) | D |  |
| Total PCR volume prepared | D | ✓ page 10 |
| Partition number | **E** | ✓ SI dPCR file |
| Individual partition volume | **E** | ✓ SI dPCR file |
| Total volume of the partitions measured (effective reaction size) | **E** | ✓ SI dPCR file |
| Partition volume variance/SD | D | ✓ SI dPCR file |
| Comprehensive details and appropriate use of controls | **E** | ✓ page 10 |
| Manufacturer of dPCR instrument | **E** | ✓ page 10 |
| **dPCR VALIDATION** |  |  |
| Optimisation data for the assay | D |  |
| Specificity (when measuring rare mutations, pathogen sequences etc) | **E** | NA |
| Limit of detection of calibration control | D |  |
| If multiplexing, comparison with singleplex assays | **E** | NA |
| **DATA ANALYSIS** |  |  |
| Mean copies per partition (λ or equivalent) | **E** | ✓ SI dPCR file |
| dPCR analysis program (source, version) | **E** | ✓ page 10 |
| Outlier identification and disposition | **E** | ✓ page 10 |
| Results of NTCs | **E** | ✓ SI dPCR file |
| Examples of positive(s) and negative experimental results as supplemental data | **E** | ✓ SI dPCR file |
| Where appropriate, justification of number and choice of reference genes | **E** | ✓ SI P23 |
| Where appropriate, description of normalization method | **E** | ✓ SI |
| Number and concordance of biological replicates | D |  |
| Number and stage (RT or qPCR) of technical replicates | **E** | NA |
| Repeatability (intra-assay variation) | **E** | NA |
| Reproducibility (inter-assay/user/lab etc variation) | D |  |
| Experimental variance or CI **^d^** | **E** | ✓ SI dPCR file |
| Statistical methods for analysis | **E** | ✓ page 10, 11 |
| Data submission using RDML (Real-time PCR Data Markup Language) | D |  |

NA: Not applicable
