## Supplementary material for "Identification of reduced host transcriptomic signatures for tuberculosis and digital PCR-based validation and quantification": STARDchecklist

|  | **Section & Topic** | **No** | **Item** | **Reported on page #** |
| --- | --- | --- | --- | --- |
|  | **TITLE OR ABSTRACT** |  |  |  |
|  |  | **1** | Identification as a study of diagnostic accuracy using at least one measure of accuracy  (such as sensitivity, specificity, predictive values, or AUC) | 1, 3 |
|  | **ABSTRACT** |  |  |  |
|  |  | **2** | Structured summary of study design, methods, results, and conclusions  (for specific guidance, see STARD for Abstracts) | 3 |
|  | **INTRODUCTION** |  |  |  |
|  |  | **3** | Scientific and clinical background, including the intended use and clinical role of the index test | 5, 6, 7 |
|  |  | **4** | Study objectives and hypotheses | 7 |
|  | **METHODS** |  |  |  |
|  | *Study design* | **5** | Whether data collection was planned before the index test and reference standard  were performed (prospective study) or after (retrospective study) | 8 |
|  | *Participants* | **6** | Eligibility criteria | 8, 9 |
|  |  | **7** | On what basis potentially eligible participants were identified  (such as symptoms, results from previous tests, inclusion in registry) | 8, 9 |
|  |  | **8** | Where and when potentially eligible participants were identified (setting, location and dates) | 8, 9 |
|  |  | **9** | Whether participants formed a consecutive, random or convenience series | 8, 9 |
|  | *Test methods* | **10a** | Index test, in sufficient detail to allow replication | 8, 9 |
|  |  | **10b** | Reference standard, in sufficient detail to allow replication | 9, 10, 11 |
|  |  | **11** | Rationale for choosing the reference standard (if alternatives exist) | 8, 9 |
|  |  | **12a** | Definition of and rationale for test positivity cut-offs or result categories  of the index test, distinguishing pre-specified from exploratory | 8,9 |
|  |  | **12b** | Definition of and rationale for test positivity cut-offs or result categories  of the reference standard, distinguishing pre-specified from exploratory | 8, 9 |
|  |  | **13a** | Whether clinical information and reference standard results were available to the performers/ readers of the index test | 8,9 |
|  |  | **13b** | Whether clinical information and index test results were available to the assessors of the reference standard | 8,9 |
|  | *Analysis* | **14** | Methods for estimating or comparing measures of diagnostic accuracy | 8, 10, 11 |
|  |  | **15** | How indeterminate index test or reference standard results were handled | 8,9,10 |
|  |  | **16** | How missing data on the index test and reference standard were handled | 8,9 |
|  |  | **17** | Any analyses of variability in diagnostic accuracy, distinguishing pre-specified from exploratory |  |
|  |  | **18** | Intended sample size and how it was determined | 8, 9 |
|  | **RESULTS** |  |  |  |
|  | *Participants* | **19** | Flow of participants, using a diagram | 12 |
|  |  | **20** | Baseline demographic and clinical characteristics of participants | 14 |
|  |  | **21a** | Distribution of severity of disease in those with the target condition | NA |
|  |  | **21b** | Distribution of alternative diagnoses in those without the target condition | Table S2 |
|  |  | **22** | Time interval and any clinical interventions between index test and reference standard | 8,9 |
|  | *Test results* | **23** | Cross tabulation of the index test results (or their distribution)  by the results of the reference standard | Table S5 |
|  |  | **24** | Estimates of diagnostic accuracy and their precision (such as 95% confidence intervals) | 11,12,13 |
|  |  | **25** | Any adverse events from performing the index test or the reference standard | NA |
|  | **DISCUSSION** |  |  |  |
|  |  | **26** | Study limitations, including sources of potential bias, statistical uncertainty, and generalisability | 24 |
|  |  | **27** | Implications for practice, including the intended use and clinical role of the index test | 23,24,25 |
|  | **OTHER INFORMATION** |  |  |  |
|  |  | **28** | Registration number and name of registry | NA |
|  |  | **29** | Where the full study protocol can be accessed | NA |
|  |  | **30** | Sources of funding and other support; role of funders | 25 |

.
